# Hierarchically-structured metalloprotein composite coatings biofabricated from co-existing condensed liquid phases

**DOI:** 10.1101/837146

**Authors:** Franziska Jehle, Elena Macías-Sánchez, Peter Fratzl, Luca Bertinetti, Matthew J. Harrington

## Abstract

Complex hierarchical structure governs emergent properties in soft biopolymeric materials; yet, the material processing involved remains poorly understood. Here, we investigated the multiscale structure and composition of the mussel byssus cuticle before, during and after formation to gain further insight into the processing of this hard, yet extensible metal cross-linked protein composite. Our findings reveal that the granular substructure crucial to the cuticle’s function as a wear-resistant coating of an extensible polymer fiber is pre-organized in condensed liquid phase secretory vesicles. These are separated into catechol-rich proto-granules enveloped in a sulfur-rich proto-matrix which fuses during secretion, forming the sub-structure of the cuticle. Metal ions are added subsequently in a site-specific way, with Fe contained in the sulfur-rich matrix and V being relegated to the granules, coordinated by catechol. We posit that this hierarchical structure self-organizes via phase separation of specific amphiphilic protein components within secretory vesicles, resulting in a meso-scale structuring, critical to the cuticle’s advanced function.

---

The ability to endow soft polymeric materials with nano- and meso-scale structural hierarchy via self-assembly is an important materials design challenge with implications for tissue engineering, drug delivery and smart polymer engineering^1–4^. Polymer scientists aim to achieve control of multiscale organization through precisely defined chemical structure and engineered supramolecular interactions^2,5,6^, which can be employed for example, as scaffolds for guiding polymerization of materials with defined hierarchy (e.g. mesoporous silica)^7–9^. Similarly, proteins also possess an inherent capacity for supramolecular self-organization – determined by amino acid sequence – that has been harnessed through evolution for fabricating bulk materials/tissues with enhanced function, controlled through multiscale hierarchical structure ^10–15^. Therefore, elucidating the physical and chemical underpinnings of biological material assembly provides a rich source of inspiration for designing bottom-up processes to fabricate hierarchically structured, functional soft materials.

Here, we report an in-depth compositional and ultrastructural investigation of the structure and self-assembly of the mussel byssus cuticle. Mussels fabricate a byssus as an attachment holdfast for anchoring on rocky surfaces in seashore habitats (Fig. 1a), where they face forces from crashing waves^16^. Each byssal thread is covered by a thin protective cuticle (Fig. 1), which is highly extensible, yet stiff and hard^17–21^. These typically opposing properties likely function to protect the stretchy fibrous core within and originate from the composite-like structure of micron-sized inclusions known as granules embedded in a continuous matrix^18^(Fig 1f-h). While it was originally proposed that the granules are hard inclusions in a softer matrix^17,18^, more recent results suggest that granules function to retain water at low hydration, behaving softer than the matrix under dry conditions^21^. Regardless, nanoindentation experiments indicate that the cuticle acquires nearly 85% of its stiffness/hardness from metal coordination cross-links between metal ions (such as Fe and V) and 3,4-dihydroxyphenylalanine (DOPA), a post-translational modification of Tyrosine elevated in the cuticle protein, mussel foot protein 1 (mfp-1) (Fig. 1i)^20^. Notably, the DOPA-metal interactions are more concentrated in the granules than the matrix^17^. (*n.b.* species specific forms of mfp-1 are named according to the species; e.g. mefp-1 for *Mytilus edulis)*

**Figure 1.**
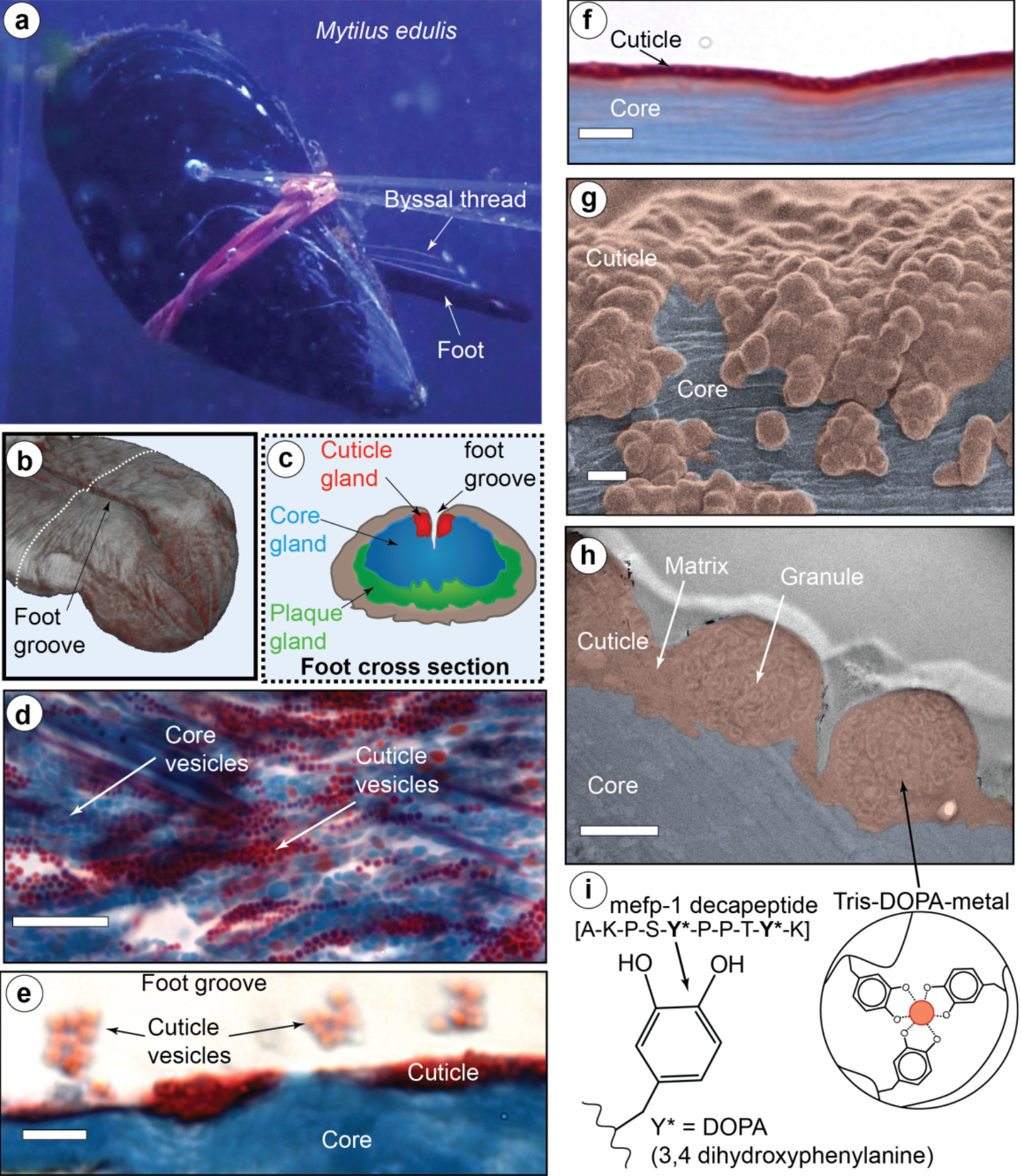
Overview of byssus cuticle formation and structure. a) Marine mussels (*Mytilus edulis*) synthesize byssal threads using a tongue-like organ known as the foot. b) CT image of a mussel foot highlighting the foot groove, in which the thread forms. c) Schematic of a foot transverse cross-section showing location of specific glands in which thread-forming proteins are stockpiled. d) Trichrome stained histological section of foot gland tissue showing the core (blue) and cuticle (red) vesicles. Scale bar = 10 μm. e) Trichrome stained thread during induced formation showing core and cuticle. Clusters of cuticle vesicles coalesce and spread over the core surface creating the cuticle. Scale bar = 4 μm. f) Trichrome stained section of a native byssal thread. Scale bar = 4 μm. g) SEM image of a native thread surface with false coloring to show the cuticle (red) and exposed core (blue). Scale bar = 1 μm. h) TEM image of a thin osmium stained section of a byssal thread with false coloring to show the cuticle and fibrous core. Scale bar = 500 nm. i) The cuticle is known to be partially comprised of a protein called mefp-1, with an extended domain made of decapeptide repeats containing 3,4-dihydroxyphenylalanine (DOPA), which is known to be coordinated to metal ions including vanadium and iron. Panels b, d, e and f are adapted from reference 13 under the Creative Commons License.

Numerous reviews highlight recent efforts to mimic the DOPA-metal cross-link strategy with mussel-inspired catechol-functionalized polymers^22–25^; yet, none of these materials reproduces the complex hierarchical structure of the cuticle or its properties. Indeed, little is understood about how the nanostructure of the cuticle and granules is achieved. It is known that the proteins that comprise the cuticle are stored within vesicles in the mussel foot – the organ responsible for synthesizing and stockpiling the proteins that make the byssus (Fig. 1b-c)^13,26,27^. Self-assembly of the cuticle occurs within minutes via release of vesicle contents into a groove running along the foot, during which they coalesce and spread over the already formed collagenous core of the byssus fiber (Fig. 1b-e)^13^. However, crucial questions remain concerning how the intricate cuticle substructure emerges via self-assembly and how metal ions are infiltrated to cure the formed cuticle. Here, we investigated the 3D nanostructure and elemental composition of the secretory vesicles and the cuticle itself, utilizing a combination of focused ion beam scanning electron microscopy (FIB-SEM), transmission electron microscopy (TEM) and scanning transmission electron microscopy with energy dispersive X-ray spectroscopy (STEM-EDS). By examining the different stages of assembly, we gain crucial new insights into the formation process and function of this complex biological material, with bearing on the design of technically and biomedically relevant composite materials.

Previous investigations have identified that the precursor proteins forming the cuticle are stored in secretory vesicles in the mussel foot within a region known as the enzyme or cuticle gland (Fig. 1d)^13,26^. TEM imaging of post-stained thin sections of the cuticle gland of *Mytilus edulis* revealed that each micron-sized vesicle possesses at least two distinctive regions – a darker outer phase (more heavily stained with Os) and a lighter, less-stained inner phase (Fig. 2a). The distinctive biphasic brain-like texture of the inner phase consisting of lighter stained connected layers was previously observed in vesicles of another species, *Mytilus galloprovincialis*^26^, and is highly reminiscent of native thread cuticle granule structure (Fig. 1h) – leading us to name this the proto-granule and the outer phase, the proto-matrix. A third, very lightly stained crescent-shaped phase is also observed at the outer periphery of most vesicles. FIB-SEM of chemically fixed foot tissue samples enabled 3D rendering of vesicles in a small region of the gland close to the secretion point, revealing a tightly packed arrangement of nearly spherical vesicles (Fig. 2b-c). Consistent with TEM, there are at least three discernible phases in the vesicles that stain differently, corresponding to the proto-granule, proto-matrix and crescent phase (Fig. 2b-d). Volumetric analysis of the 3D data of 28 vesicles indicates similar distribution of volume fractions of the three phases in all vesicles suggesting a highly regulated formation process (Fig. S1).

**Figure 2.**
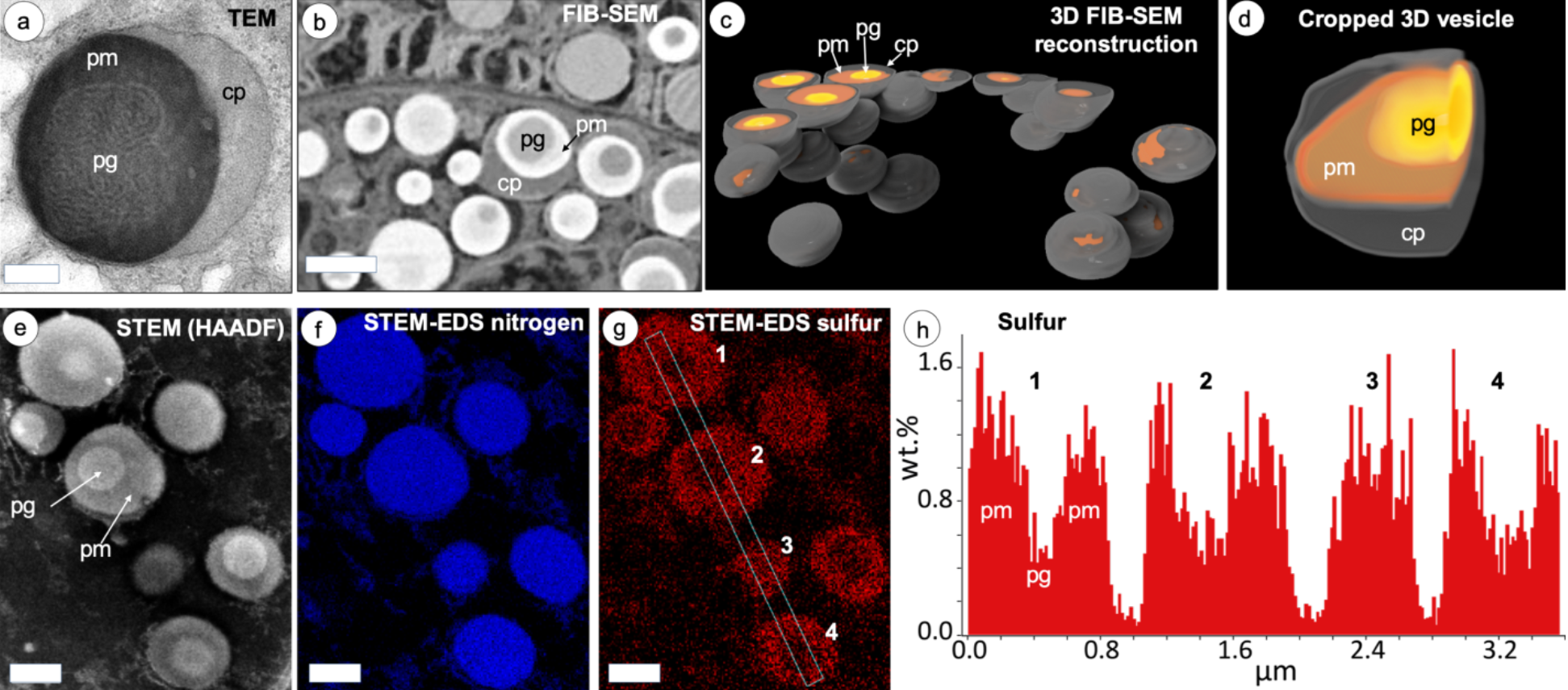
TEM and FIB-SEM imaging of cuticle secretory vesicles in mussel foot tissue. a) TEM image of osmium stained cuticle secretory vesicle from region similar to Fig. 1d. Contrast is achieved by different osmium staining with the proto-matrix (pm) staining the heaviest, the crescent phase (cp) staining the lightest and the proto-granules (pg) staining in between. Scale bar = 200 nm. b) FIB-SEM image of osmium stained region of cuticle gland similar to Fig. 1d. Contrast is inverted compared to TEM, but pg, pm and cp can clearly be differentiated. Scale bar = 1000 nm. c) Using electron density contrast, secretory vesicles and their inner structure were reconstructed in 3D from an image stack consisting of 392 images. d) Magnified and cropped 3D image of a vesicle showing all three phases. e) HAADF image (STEM mode) showing localization of pm and pg phases in vesicles consistent with TEM and FIB-SEM. f) STEM-EDS of nitrogen signal is uniform between pg and pm. Scale bar for panels e-g = 500 nm. g) In contrast, STEM-EDS of sulfur shows differences. h) Relative sulfur wt% (not calibrated) collected from 4 vesicles in dotted box in panel (g) reveals that pg has approximately half the amount of sulfur at pm.

Compositional analysis of vesicles was performed with STEM-EDS, revealing an elevated and uniform nitrogen content in vesicles compared with the surrounding intracellular material (Fig. 2e-f). In contrast, there is an approximately two-fold higher sulfur content in the proto-matrix relative to the proto-granules (Fig. 2g-h). Notably, recent transcriptomic studies have identified a family of putative cysteine-rich proteins (mfp-16 – mfp-19) within the cuticle gland of a related species *Mytilus californianus*^28^. This is consistent with previous cytochemical studies predicting the presence of cysteine-rich proteins within the secretory vesicles of *Mytilus galloprovinicialis*^26^. The fact that the proto-granule contains a lower sulfur signal substantiates the previous supposition that the granules mainly contain mefp-1, which completely lacks Cysteine^17^. Furthermore, Cysteine is a strong target of OsO4 staining^29,30^, which helps explain the contrast observed in TEM and FIB-SEM. Notably absent in the vesicles are transition metal ions (e.g. Fe and V) that are present in the mature cuticle coordinated to DOPA^17,19,20^. This result is consistent with recent findings suggesting that metal ions are added into the cuticle in a post-secretion curing step^13^.

To investigate the dynamic transition from the storage phase of vesicles into a hard and flexible mesostructured coating, TEM was performed on vesicles secreted into the foot groove. Vesicle secretion was induced by injection of 0.56 KCl into the pedal nerve of the foot, as previously used to study thread formation^13,31^. TEM images of induced threads chemically fixed several minutes into the formation process reveal that cuticle vesicle contents fuse and coalesce already within the secretory ducts leading from the glands to the foot groove (Fig. 3a). This reveals that the proto-matrix retains a liquid-like behavior as it coalesces with the contents of other vesicles, showing no apparent boundaries. However, the proto-granules remain as separate entities with the biphasic brainy structure intact. Within the groove, the coalesced vesicles spread over the collagenous core forming a thin layer, as previously shown with histology and confocal Raman spectroscopy (Fig. 1e and 3b-c)^13^. This granular morphology is very similar to the native cuticle structure (Fig. 1h). Taken together with the TEM-EDS measurements (Fig. 2e-g), these observations suggest that the vesicles contain two immiscible protein phases represented by the proto-matrix and proto-granule that are pre-organized within vesicles to facilitate rapid assembly of a hierarchically structured coating.

**Figure 3.**
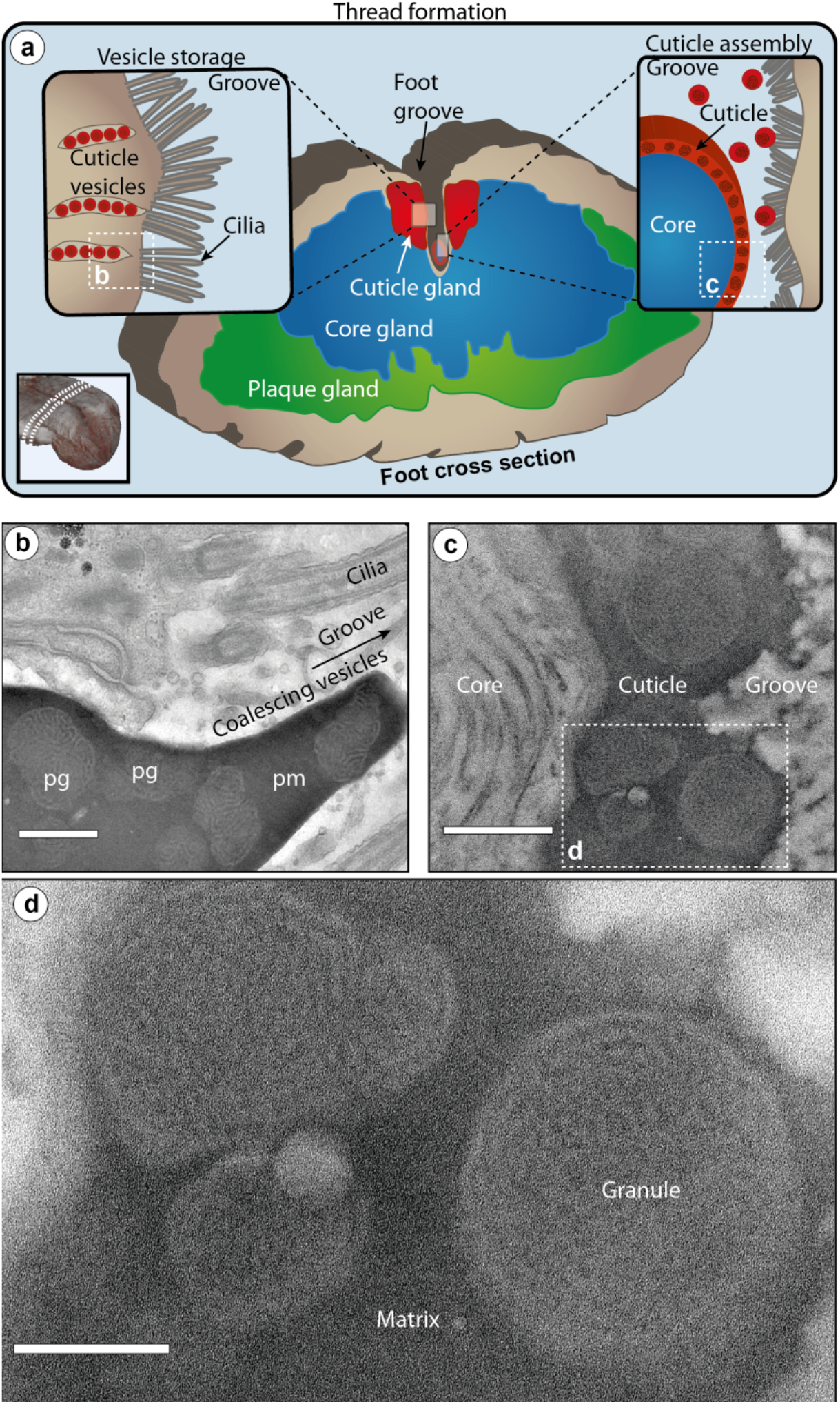
Cuticle formation in the mussel foot groove via coalescence of vesicle contents. a) Schematic overview of transverse section of mussel foot tissue showing anatomy of byssal glands. Cuticle vesicles are lined up at the edge of the cuticle gland, ready to be secreted into the groove, aided by cilia. During cuticle assembly, the vesicle contents cluster and coalesce into the groove spreading over the formed core surface. b) TEM image from stained section of an induced foot in the region indicated in panel (a), showing coalesced cuticle vesicles about to be released into the groove. Notably, the proto-matrix (pm) of several vesicles merges to form a continuous matrix, while the proto-granules (pg) retain their brain-like structure. Scale bar = 500 nm. c) TEM image from stained section of an induced foot in the region indicated in panel (a), showing a formed induced thread with intact cuticle. Scale bar = 500 nm. d) Magnified region from panel (c) showing details of the induced cuticle including granule and matrix. Scale bar = 200 nm.

The nanoscale structural and compositional details of the native cuticle were next investigated with FIB-SEM and STEM-EDS on chemically-fixed thread samples, both of which revealed that *M. edulis* thread cuticles consist of a single layer of granules, rather than the several layers observed in other *Mytilus* species^18,21^. FIB-SEM analysis enabled the visualization of the matrix and granules, as well as the convoluted brain-like internal substructure of the granules (Fig. 4a-b). 3D reconstruction of the image stacks revealed the connectivity of the matrix phase and heavily stained part of the granule in three dimensions and further indicates that the lightly stained (darker) region within the granules is comprised of a connected network of flattened layers resembling a bicontinuous phase of a micro phase-separated block co-polymer (Fig. 4c-d) ^32^. Image analysis reveals the flattened layers of multiple granules within a given thread possess a highly-defined thickness of ~20 nm, although this varied between different threads (Fig. 4d, S2, Table S1). Additionally, the flattened layers within the granules are oriented along a common direction approximately 45° to the fiber axis (Fig. 4c) – the orientation of which is consistent between different granules within the same thread. The 3-dimensional (spherically averaged) autocorrelation functions of the different measured granules are similar to each other within a given thread, indicating a high homogeneity in their structural features (Fig. S2).

**Figure 4.**
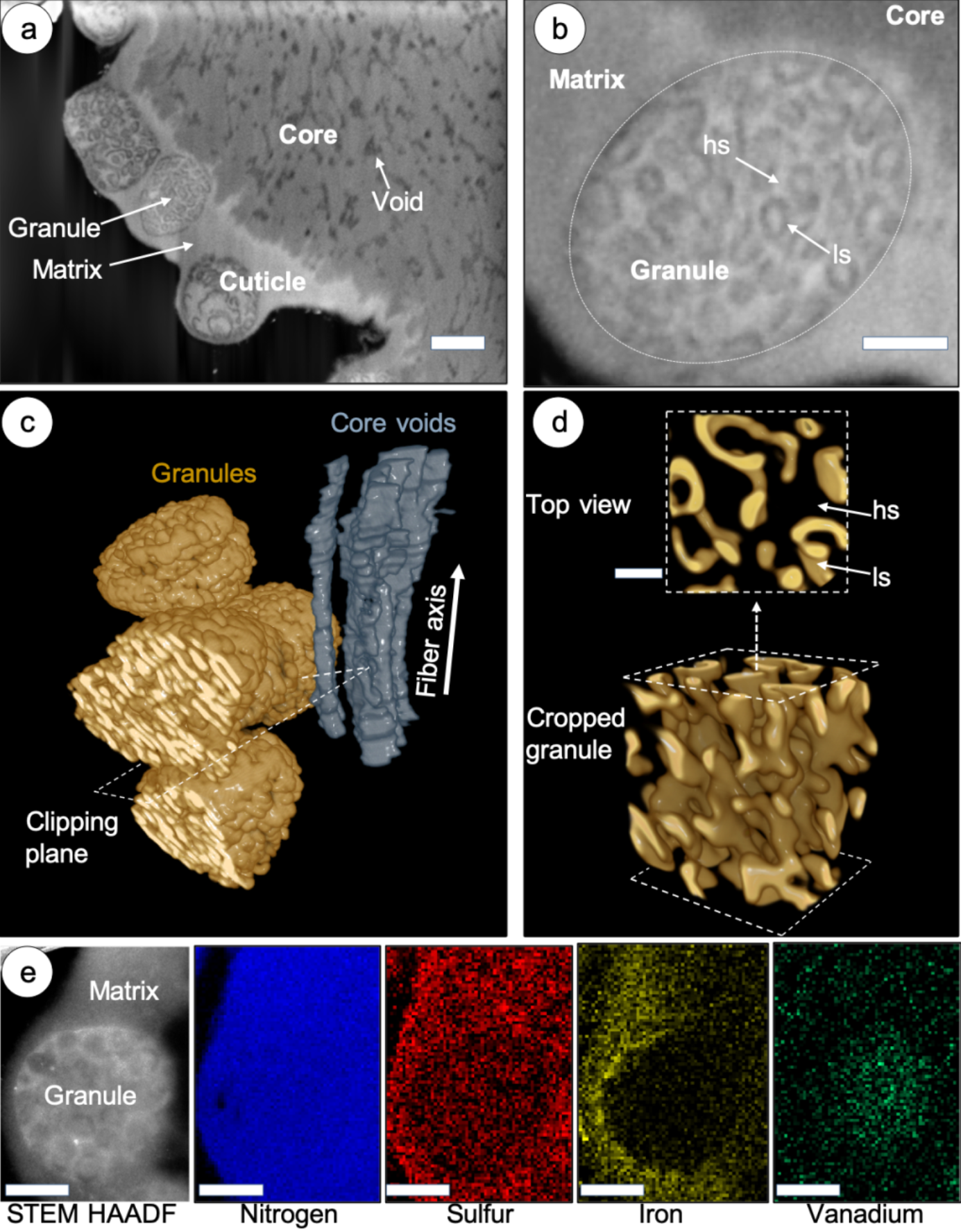
FIB-SEM 3D-reconstruction of intragranular nanostructure. a) FIB-SEM image of osmium stained region of native thread showing substructure of the core and cuticle, including granule and matrix in the cuticle and previously described voids in the core. Scale bar = 500 nm. b) Higher magnification FIB-SEM image of a single granule. Contrast arises from heavily staining (hs) and lightly staining (ls) regions within a single granule. *n.b.* contrast is inverted compared to TEM with higher Z elements appearing brighter. Scale bar = 200 nm. c) 3D reconstruction of FIB-SEM image stack using the contrast of the lightly staining (ls) phase of the granules and the core voids (which are parallel to the fiber axis). d) Granule cropped using clipping planes parallel to the one indicated in (c) revealing internal structure of the ls phase, consisting of bicontinuous flattened layers with a thickness of ~20 nm. The transparent region between layers constitutes the heavily staining (hs) phase from panel (b). Scale bar = 80 nm. e) STEM-EDS compositional analysis of native cuticle granule and matrix regions, showing distribution of nitrogen, sulfur, iron and vanadium in the region in the STEM-HAADF image on the left. Scale bar = 100 nm.

At first glance, STEM-EDS compositional studies of the mature cuticle are largely consistent with those measured on the secretory vesicles. Similar to the proto-granules and proto-matrix in the vesicles, the granules and matrix have uniform nitrogen levels, while the matrix phase has a significantly higher sulfur signal than the granules (Fig. 4e, S3). However, while transition metal ions were not detected in the secretory vesicles, both Fe and V were detected by EDS in the native cuticle (Fig. 4e). Unexpectedly, however, metal distribution was micro-partitioned within the cuticle, with V explicitly associated with the granules and Fe associated only with the matrix with a sharp interface at the boundary. This is consistent with previous Raman spectroscopic measurements across different *Mytilid* species showing that DOPA-V coordination is predominantly detected in native cuticles, rather than DOPA-Fe coordination – although it should be noted that Fe can be artificially introduced and coordinated following removal of V with EDTA^17,20^. Because the metals are not present in the secretory vesicles, this finding also implies that the metal ions spontaneously segregate between the matrix and granules when introduced during thread formation.

The findings of this study suggest that the granular mesostructure of the cuticle is achieved through a membrane-bound liquid-liquid phase separation (LLPS) consisting of immiscible fluid protein phases. Co-existing condensed liquid protein phases are observed in nucleolar subcompartments and believed to be important for tuning the vectorial transport and processing of rRNA^33^. In contrast, within the cuticle vesicles, the co-existing LLPS leads to a micro-to nanoscale distribution of proteins with different functional groups (i.e. DOPA and Cysteine) that contribute to cross-linking the liquid phase to solid when triggered during assembly. This effectively tunes the viscoelastic properties of the final material at the mesoscale. Although deducing a biologically evolved function *ex post facto* is tricky, we posit that microscale partitioning of specific metal coordination cross-links between the granules and matrix has crucial implications for the dynamic properties of this material, and it is important to understand why and how this occurs. Current models postulate different mechanical properties in the matrix and granules – either hard granular inclusions in a soft matrix^17,18^ or soft, water-absorbent granules in a stiffer matrix (at least under low hydration conditions)^21^. However, both models rest on results from quasistatic mechanical indentation performed at a single loading rate, which do not access the dynamic nature of the metal coordination bonds. On the other hand, rheological investigations of mussel-inspired catechol-enriched metallopolymers have demonstrated clearly that DOPA-V coordination bonds possess a more than 10-fold longer bond lifetime (and thus, slower relaxation time) than DOPA-Fe bonds^34^. This is functionally relevant considering that mussels face an enormous range of mechanical loading rates from rapid crashing waves with velocities up to 30 m/s to the extremely slow loading from predators such as sea stars^16^. Thus, it seems plausible that the physical demands of life in the intertidal have led to evolution of an adaptive coating that responds to a wide range of strain rates, achieved through hierarchical organization of dynamic bonds.

Fabricating this functional composite structure reproducibly requires a remarkable degree of process control during assembly achieved by pre-organizing the various cuticle components into phase separated LLPS within the secretory vesicles prior to assembly (Fig. 2). Until recently, only DOPA-rich mefp-1 was confirmed to be present in the cuticle (Fig. 1i). However, as mentioned, transcriptomics has identified four new putative cuticle proteins (mfp-16 – mfp-19) in a closely related species^28^, all of which are enriched in Cysteine and likely present primarily in the proto-matrix/matrix, based on our STEM-EDS findings and previous cytochemical evidence^26^. The rapid coalescence of the proto-matrix into a solid material during formation is likely related to the well-known ability of Cysteine to participate in various covalent and non-covalent interactions^35^. However, as previously proposed, Cysteine in byssus proteins likely also perform a crucial role in redox cycling of the DOPA catechol moiety^36^. Both putative roles will be discussed later in more detail. Evidence that mefp-1 is localized in the granule comes from the previous Raman-based localization of DOPA-V interactions in the granules^17,20^, the different susceptibilities of the granule to chymotrypsin vs. pepsin digestion in cytochemical studies^26^ (Fig. S4) and the lower sulfur content in the granules observed here with STEM-EDS (Fig. 2g and 4e).

Given these compositional findings, what is then the source of complex nano-structure observed in the secretory vesicles, and thus, the final cuticle structure? We propose that the bicontinuous structure and lower sulfur signal of granules arises from the immiscibility of mefp-1 and the Cys-rich protein, which is driven by the amphiphilic block co-polymer-like structure of mfp-1 (Fig. S5). Mefp-1 consists of numerous repeats of the decapeptide consensus motif [AKPS<u>Y</u>PPT<u>Y</u>K]n in which the tyrosine (Y) residues are largely converted to DOPA with a total content of 10 – 15 mol% and known to interact with metal ions (Fe, V) (Fig. 1i, 5, S5)^17,20^. The repetitive region of mefp-1 possesses a hydrophilic character due to the high content of conserved Lysine (K) residues. Recombinant expression of truncated mfp-1 consisting of 12 or 22 repeats of the hydrophilic decapeptide have been demonstrated to undergo spontaneous LLPS under high salt conditions mediated via pi-cation interactions^37^, offering support to the coacervate-like nature in the vesicles. However, at the N-terminus of native mefp-1 and mgfp-1 from *M. galloprovincialis* is an 60-80 amino acid non-repetitive domain that is markedly less hydrophilic, giving the overall protein an amphiphilic profile (i.e. block co-polymer-like) (Fig. 5, S5). We propose here that the non-repetitive domain provides the impetus for formation of a bicontinuous phase (also observed in *M. galloprovincialis),* forming the well-defined ~20 nm thick layers characteristic of the granules, while the positively charged DOPA-rich phase interacts with the Cysteine-rich proteins to produce the heavily stained phase of variable thickness between the flattened layers (Fig. 5). The fact that osmium has a very high affinity to Cysteine under the alkaline staining conditions supports this model^29,30^. The matrix on the other hand, seems to consist mostly of the Cysteine-rich proteins.

**Figure 5.**
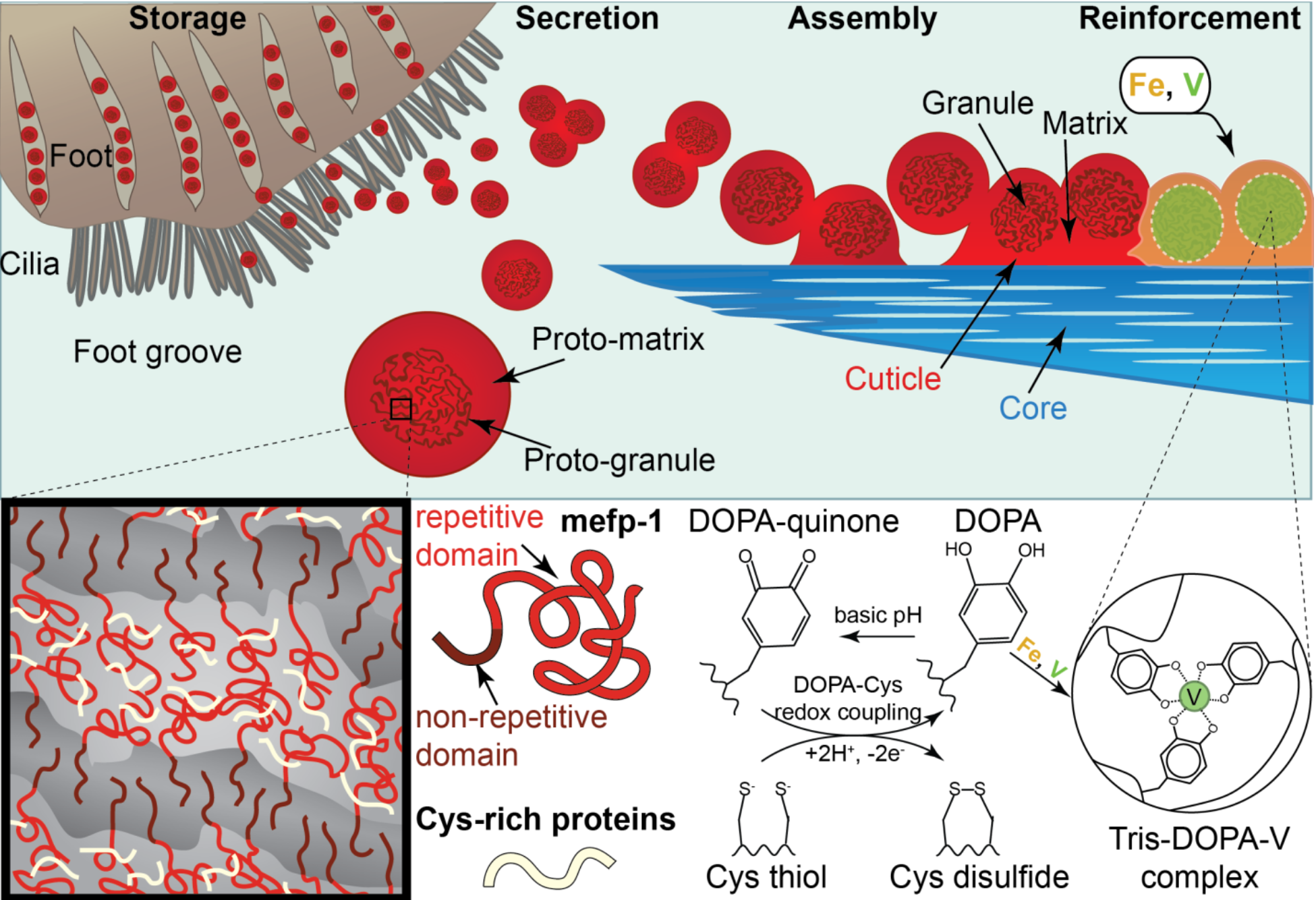
Schematic model of cuticle assembly via co-existing condensed liquid phase vesicles. To achieve the high degree of compositional, structural and mechanical hierarchy observed in the native byssus cuticle, mussels store the protein precursors as a LLPS of two co-existing phases. The main protein component mfp-1 and the Cys-rich proteins (mfp-16-19) are immiscible due to the amphiphilic nature of mfp-1, leading to phase separation into a bicontinuous structure that characterizes the proto-granule. During secretion and cuticle assembly, the proto-matrix of nearby vesicles fuses forming the continuous matrix of the cuticle, possibly cross-linked via Cysteine residues. The intricate nanostructure of the proto-granule is maintained in the newly formed cuticle and contains a much higher local concentration of DOPA than the surrounding matrix. Thus, when metals are added to the thread in a secondary curing step (mechanism still not understood), the metal ions that have the highest stability complex with DOPA (i.e. vanadium) diffuse into the granule where they become concentrated relative to iron. This results in different viscoelastic behavior of the granule and matrix, which is likely an adaptive function to life on the highly dynamic rocky seashore. Cysteine may play a secondary, but equally important role as a reducing agent that counteracts the spontaneous oxidation of DOPA to DOPAquinone under basic conditions, which enables strong metal coordination cross-link formation.

Based on this new model of how the observed meso- and nano-structures form in the vesicles, we consider next how the condensed fluid phase of the vesicles suddenly transitions into a hard, yet stretchy composite material. Our TEM investigation of induced thread formation indicates that the Cysteine-rich proto-matrix fuses during assembly, creating the continuous matrix of the cuticle in which the pre-assembled granules are embedded. As mentioned, Cys is a highly promiscuous cross-linker with the ability to form covalent bonds via disulfide linkages^35^ or through oxidative cross-linking with DOPA residues^36^. In fact, 5-cysteinyl-DOPA was previously purified from byssus material^38^. However, considering that nearly 85% of cross-links in the cuticle depends on metal ions^20^, Cysteine-based covalent bonds are not the dominant cross-linking mechanism in the matrix. Based on the co-localization of Cysteine and Fe in the matrix, we consider the compelling possibility that sulfur-Fe clusters may contribute as load-bearing crosslinks. While typically found in enzymes involved with redox pathways in cells, recent AFM single molecule force spectroscopic reveal that Fe-S clusters can function as reversible sacrificial crosslinks in proteins with mechanical breaking force comparable to other metal coordination complexes^39^.

A secondary, but equally important role of the Cysteine residues within the granules may be as a reducing agent to stall spontaneous DOPA oxidation in the basic environment of the ocean (Fig. 5). Another Cysteine-rich protein called mfp-6 was previously identified in the adhesive secretion of the byssus^38^ and was shown *in vitro* prevent DOPA from spontaneously oxidizing to DOPAquinone (a less effective adherent) through redox cycling^36^. Similarly, DOPA-quinone is inefficient at forming strong metal coordination complexes^40^. Considering that at least 85% of cross-linking in the cuticle is based on metal coordination bonds^20^, the proposed reducing role of Cysteine may be crucial to achieving the dynamic properties, and is likely enhanced by the proposed colocalization with the DOPA-rich repetitive domain of mefp-1 in the granules (Fig. 5).

Shortly after the cuticle forms during assembly via fusion of the proto-matrix, transition metal ions (Fe and V) are apparently added, providing secondary mechanical reinforcement via metal coordination bonding^13,17,20^. Raman spectroscopic investigation of the cuticle revealed that DOPA is able to bind both V and Fe within the cuticle when exposed to only the individual ions during *in vitro* experiments^17,20^. However, all Raman spectra reported for native byssal threads across several species are dominated by the DOPA-V signal, rather than DOPA-Fe^17,20^. It is tempting to propose that this partitioning may reflect the inherent differences in dissociation rates of the tris-DOPA-Fe and tris-DOPA-V complexes previously revealed by rheological studies of mussel-inspired catechol-functionalized hydrogels showing that DOPA-V has more than an order of magnitude longer bond lifetime than DOPA-Fe complexes^34^. The slower bond dissociation of DOPA-V implies that it will be less likely to exchange with Fe once coordinated, leading to its concentration in the mfp-1-enriched granules over time through dissociation and diffusion. However, the longer bond lifetime, not only affects the distribution of metals, but as already pointed out, will also determine local viscoelastic relaxation behavior defined by the compartmentalization of mfp-1 within the granules. Taking for granted that varying the viscoelastic response on the mesoscale is important to the function of the cuticle (e.g. being hard and extensible, retaining water in dry conditions and/or responding to a range of loading rates^17,18,20,21^), this assembly mechanism provides a degree of control over dynamic mechanical response that is not possible in current engineered polymers. Indeed, mimicking this fabrication process and thus, this remarkable degree of control over soft matter response could inspire the design of new responsive smart polymers for a range of applications from flexible electronics and actuated structures to drug delivery and dynamic tissue scaffolds.

## Methods

### Materials

Blue mussels (*M. edulis*) purchased from the Alfred-Wegener-Institut were maintained at ~ 14 °C in an aquarium with artificial salt water. Investigations were performed either on byssal threads or on the foot organ of adult mussels (5 – 8 cm in length) removed with a scalpel. To investigate cuticle assembly, protein secretion was induced by injecting 0.56 M KCl solution in the base of two mussel feet as previously described^13,31^. Induced mussel feet were dissected after at ~5 min following KCl injection to investigate vesicle secretion and after ~20 min to investigate the induced thread cuticle structure. Collected native threads were stored at 4 °C in water prior to use.

### Chemical fixation and embedding

Dissected feet (induced or not) were carefully rinsed with cold water, blotted with a paper towel to remove mucus and pre-fixed for 30 min at 4 °C in 3 % glutaraldehyde, 1.5 % paraformaldehyde, 650 mM sucrose in 0.1 M cacodylate buffer pH 7.2. The foot tissue was then cut into thin transverse sections comprising the groove and part of the gland tissue and then fixed for 2 h at 4 °C in the same buffer as above. Fixed samples were rinsed 5× with 0.1 M cacodylate buffer, pH 7.2 at 4 °C and post-fixed with 1 % OsO4 for 1 h at 4 °C. Tissue samples prepared for elemental analysis were not treated with OsO4. Samples were rinsed again in 0.1 M cacodylate buffer pH 7.2 (3 × 5 min at 4 °C), followed by series dehydration in ethanol (50 %, 70 %, 90 %, 3 x 100 %) for 10 min each step at RT. Dehydrated samples were embedded either in low viscosity Spurr’s resin (Electron Microscopy Sciences, # 14300) for TEM/STEM-EDS or in Hard Plus resin 812 (Electron Microscopy Sciences, # 14115) for FIB-SEM and polymerized at 70 °C for at least 48 h. Ultrathin sections of 100 nm for TEM investigations were prepared using a PowerTome Model XL ultramicrotome (Boeckeler Instruments, Inc.) and mounted on carbon coated Cu grids (200 mesh) for imaging and on Lacey carbon coated Cu grids (200 mesh) for EDS measurements. In order to reveal the internal structure of the protogranule, some grids were post-stained with 2% uranyl acetate for 10 min.

The distal region of native threads were washed 3 x 5 min in cold double distilled H2O and cut into small pieces of ~3 mm in length. Fixation was carried out for 1 h at 4 °C in 2.5 % glutaraldehyde and 1.5 % paraformaldehyde in 0.1 M cacodylate buffer pH 7.4. Samples were rinsed 3 × 10 min in 0.1 M cacodylate buffer at 4 °C before post-fixation with 1 % OsO4 for 1 h. Samples for EDS measurements were not stained with OsO4. A second rinsing step in 0.1 M cacodylate buffer pH 7.4 (3 × 5 min at 4 °C) was followed by dehydration in ethanol (50 %, 70 %, 90 %, 3 × 100 %) for 10 min each step at RT. Threads were embedded in low viscosity Spurr’s resin (Electron Microscopy Sciences, # 14300) at 65 °C over two days. The resulting resin blocks were trimmed to the region of interest and sectioned to 100 nm using an ultramicrotome (PowerTome Model XL). Ultrathin sections were mounted on lacey carbon coated copper grids (200 mesh) for imaging and EDS measurements.

### Transmission Electron Microscopy

Transmission Electron Microscopy was performed with a Zeiss EM 912 Omega with an acceleration voltage of 120 kV and a Jeol JEM ARM200F equipped with a cold field-emission electron source and a Silicon Drift Detector (SSD), operated at 200 kV acceleration voltage and 15 μA emission current. TEM mode was used exclusively for imaging (Bright Field imaging) at magnifications of 10,000× and 16,000×, whereas STEM mode was used for Energy Dispersive Spectroscopy (EDS) and High Angle Annular Dark Field (HAADF) imaging. In STEM mode a fine electron probe scans the surface of the sample pixel-by-pixel enabling to identify the area of the sample that generates certain characteristic X-rays with nanometric resolution. Overview elemental maps of foot sections were acquired at a magnification of 50,000× with a pixel size of 17 nm x 17 nm and an exposure of 1 sec per pixel. For thread sections, elemental maps (59 × 89 pixel) were acquired at a magnification of 200,000× with a pixel size of 9.3 × 9.3 nm and an exposure of 1 sec per pixel using a Silicon Drift Detector (SDD).

### FIB-SEM

Resin blocks containing samples were polished in order to expose the tissue or thread at the block surface. Samples were sputter-coated with three Carbon layers (~ 5 nm each) and one platinum layer (~ 5-10 nm) and transferred to the Zeiss Crossbeam 540 (Carl Zeiss Microscopy GmbH, Germany). At the region of interest, a trench for SEM imaging was milled into the sample surface using the 65 nA FIB current at 30 kV acceleration voltage. The resulting cross-section was finely polished using the 1.5 nA FIB probe at 30 kV. Measurement of foot tissue and threads required different parameters. Thin slices of samples were removed in a serial manner by FIB milling (300 pA, 30 kV, slice thickness 17.5 nm for foot tissue; 700 pA, 30 kV, slide thickness 10.5 nm for threads). After each milling step, the specimen was imaged by SEM using the secondary and backscattered electron detector (acceleration voltage = 2 kV for foot tissue and 2.5 kV for threads). For foot tissue and threads, the image resolution was 2048 × 1536 pixels and 1024 × 785 pixels, respectively with a lateral image pixel size of 12.4 nm and 4.8 nm, respectively. Images were recorded using line averaging (N = 4 for foot tissue and N = 31 for threads) and a dwell time of 200 ns.

### FIB-SEM Data Processing

The resulting secondary and back-scattered electron images were processed using the SPYDER3 (Scientific Python Development Environment) (Python 3.6) software. Custom-written python scripts were developed and provided by Luca Bertinetti. For data analysis of cuticle gland tissue, images were automatically aligned using enhanced correlation coefficient alignment. Total variation denoising was performed by applying the Chambolle’s projection algorithm (100000 iterations, weight 0.07, 0.001 eps) in 3D mode. Segmentation of cuticle vesicles was performed using the ZIB version of Amira 3D (Thermo Fisher Scientific, USA). Vesicle shapes of 28 individual granules were segmented manually from 392 processed back-scattered electron images using the brush tool. The inner phase, which corresponds to the proto-granules as well as the proto-matrix phase were automatically segmented using the Magic Wand tool. 3D visualization of both phases was realized by volume rendering of segmented structures.

For data analysis of the thread cuticle structure, secondary electron images were automatically aligned using the Fourier shift theorem for detecting the translational shift in the frequency domain and vertical stripes arising from the water fall effect by FIB milling were removed by Fourier filtering. Total variation denoising was performed by applying the Chambolle algorithm in 3D mode. The images were inverted afterwards with Fiji and Sauvola’s local thresholding computation was applied with a block size of 11, a k value of 0.005 and r value of 1.7. As the thresholded images contained regions resulting from statistical noise in the image, only thresholded regions containing minimum 40 pixels were selected. Images were median 3D filtered with Fiji and x, y, z radii were set to 1.5. Segmentation of five adjoining cuticle granules was performed using the ZIB version of Amira 3D (Thermo Fisher Scientific, USA). Granule shapes were segmented manually from 352 processed and inverted secondary electron images using the brush tool. In a second step, the lightly staining (ls) granule phase was segmented automatically from local threshold computed and median3D filtered image stacks using the Magic Wand tool.

## Acknowledgements

The authors acknowledge funding from the Max Planck Society and the German Research Foundation (DFG individual grant HA 6369 5). M.J.H. acknowledges support from the support from the Natural Sciences and Engineering Research Council of Canada (NSERC Discovery Grant RGPIN-2018-05243).

## Author Contributions

M.J.H., L.B. and P.F. devised and oversaw the study. L.B., F.J. and E.M.-S. performed experiments and processed data. M.J.H. and F.J. wrote the paper. All authors contributed to editing the paper.

## Competing financial interests

The authors declare no competing financial interests.

