## Supplemental Materials for "Hierarchically-structured metalloprotein composite coatings biofabricated from co-existing condensed liquid phases"

### Volumetric analysis of 3D reconstructions of cuticle vesicles from FIB-SEM data

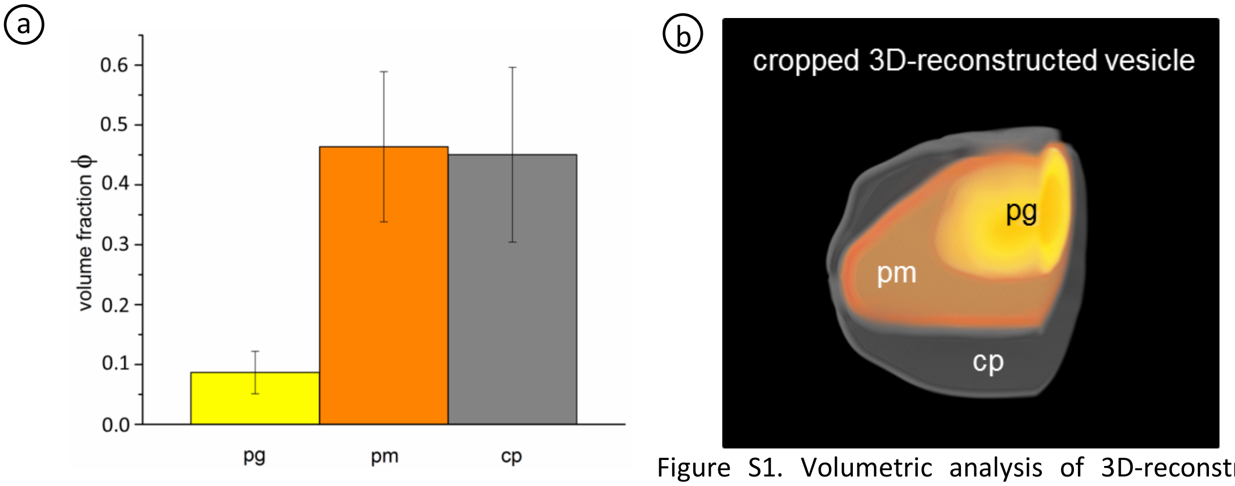


Figure S1. Volumetric analysis of 3D-reconstructed cuticle vesicles. a) Volume fractions of the proto-granule (pg), proto-matrix (pm) and crescent phases (cp) (as indicated in b). Volume analysis was performed on 28 reconstructed vesicles, showing very similar results and suggesting a highly regulated formation process.

**Analysis of granular substructure in native thread cuticle**

**Table S1. Layer thickness of granule layers extracted from FIB-SEM 3D reconstructions**

| | | Thickness Mean (nm) | Standard Deviation (nm) |
|---|---|---|---|
| **thread a** | **granule 1** | 23.63 | 5.65 |
| | **granule 2** | 23.44 | 5.32 |
| | **granule 3** | 23.44 | 5.34 |
| | **granule 4** | 24.57 | 5.66 |
| | **granule 5** | 23.71 | 5.56 |
| **thread b** | **granule 1** | 17.00 | 3.99 |
| | **granule 2** | 16.46 | 4.00 |
| | **granule 3** | 15.23 | 3.54 |
| | **granule 4** | 15.22 | 3.53 |
| | **granule 5** | 17.72 | 4.37 |

To investigate the structure of the flattened bicontinuous layer within the granules Fig. 4c-d, we analyzed the layer thickness using the BoneJ2 plugin[1] for the Fiji imaging software[2] originally programmed for calculating thickness in trabecular bone, but applicable to the granule nanostructure as well. The average thickness of the flattened layer was measured for 5 granules in each of the two different threads as shown in Table S1. The mean values for each thread were found to be 23.8 ± 5.5 nm for thread a and 16.3 ± 3.9 nm for thread b. Consistent with these measurements, Fig. S2 shows the azimuthal integration of the 3D Fourier transform of the image stacks of the same 5 granules from each of the two threads, showing a strong interthread similarity and slight differences between the threads.

ⓐ ⓑ

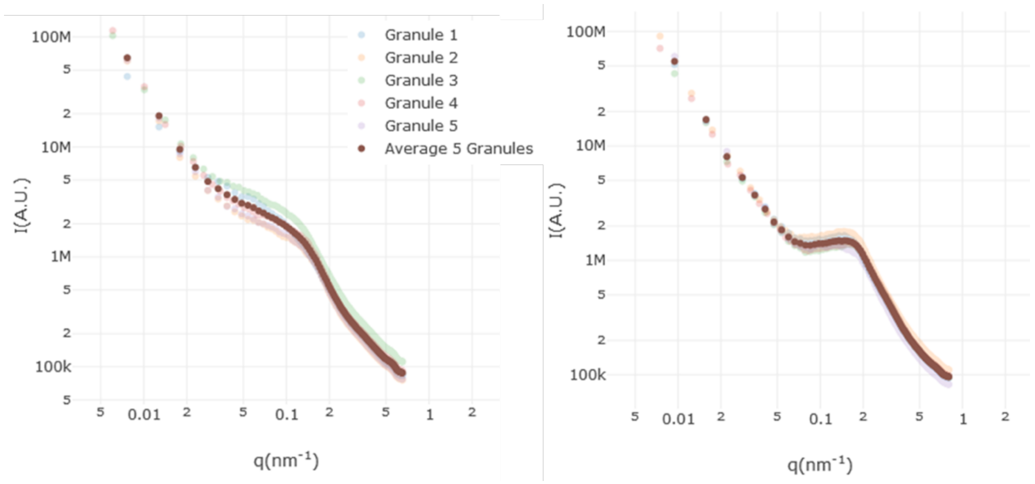


Figure S2. Azimuthal integration of the 3D Fourier transforms of the image stacks of 5 granules from each of two different investigated thread cuticles (a and b). For a given thread profiles are similar to each other, suggesting a high homogeneity in their structural features.

In spite of the clear differences between the two threads, it is remarkable how consistent the thickness between different granules within a single thread is. This suggests that the variation between the two threads might be based either on inherent differences in biological processing at various stages during the formation of the individual threads or experimental variation during sample preparation and fixation (e.g. different degrees of drying during ethanol treatment). In spite of these differences, the overwhelming similarity of the structure between granules in a single thread is highly suggestive that this assembly process is under a considerable degree of control.

**Elemental mapping of native thread cuticle with STEM-EDS**

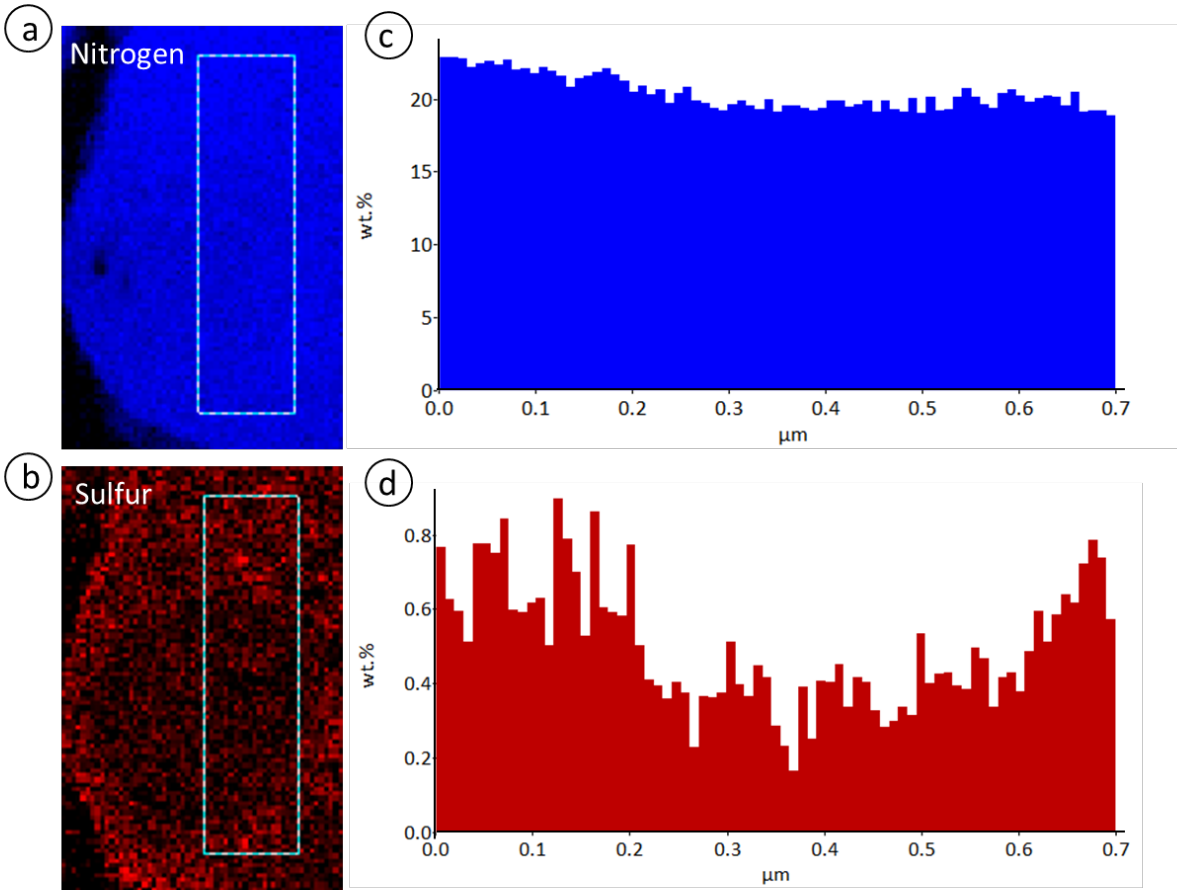


Figure S3. STEM-EDS maps showing the distribution of Nitrogen a) and Sulfur b) in cuticle matrix and granule. c) Relative nitrogen wt % (not calibrated) in dotted box (a) shows a relatively constant amount of nitrogen in the matrix and granule region. d) Relative Sulfur wt % (not calibrated) in dotted box (b) shows that sulfur is approximately two times more concentrated in the matrix than in the granule.

### **Cytochemical evidence for the presence of mfp-1 in the cuticle granules**

Previously it was shown that the granule part of the secretory vesicle from *Mytilus gallopvroncialis* (closely related to *M. edulis* and also possessing brain-like granules) is susceptible proteolytic digestion by Chymotrypsin in thin sections, but not by pepsin[3]. It was previously suggested that the granules are enriched in mfp-1[4]. In order to test whether mgfp-1, may have been the target of chymotrypsin, yet resistant to pepsin, we performed an in silico proteolytic analysis of the mgfp-1 sequence with the Peptide Cutter program (EXPASY). Indeed, Peptide Cutter predicts over 86 cleavage sites with chymotrypsin, while only 8 are predicted with pepsin, adding additional support that mfp-1 is localized in the granules.

ⓐ Amino acid sequence of mgfp-1

```
        10         20         30         40         50         60
FSNGNIYNAH GSAYAGASAG AYKTLPNAYP YGTKHGPVYK PVKTSYHPTN SYPPTYGSKT

        70         80         90        100        110        120
NYLPLAKKLS SYKPIKTTYN AKTNYPPVYK PKMTYPPTYK PKPSYPPTYK PKPSYPATYK

       130        140        150        160        170        180
SKSSYPSSYK PKKTYPPTYK PKLTYPPTYK PKPSYPPTYK PKPSYPATYK SKSSYPPSYK

       190        200        210        220        230        240
TKKTYPSSYK PKKTYPSTYK PKVSYPPTYK SKKSYPPIYK TKASYPSSYK PKKTYPSTYK

       250        260        270        280        290        300
PKISYPPTYK AKPSYPTSYR AKPSYPSTYK AKPSYPPTYK AKPSYPPTYK AKPTYPSTYK

       310        320        330        340        350        360
AKPSYPPTYK AKPSYPPTYK AKPSYPPSYK PKTTYPPSYK PKISYPPTYK AKPSYPPIYK

       370        380        390        400        410        420
AKPSYPPTYK AKPSYLPTYK AKPSYPPTYK AKPRYPTTYK AKPSYPPTYK AKPSYPPTYK

       430        440        450        460        470        480
AKLSYPPTYK AKPSYPPTYK AKPSYPPTYK AKPSYPPTYK TKPSYPRTYK AKPSYSSTYK

       490        500        510        520        530        540
AKPSYPPTYK AKPSYPPTYK AKPSYPPTYK AKPSYPPTYK AKPSYPPTYK AKPSYPQTYK

       550        560        570        580        590        600
AKSSYPPTYK AKPSYPPTYK AKPSYPPTYK AKPSYPPTYK AKPSYPPTYK AKPSYPPTYK

       610        620        630        640        650        660
AKPSYPPTYK AKPSYPPTYK AKPSYPPTYK AKPSYPATYP STYKAKPSYP PTYKAKPSYP

       670        680        690        700        710        720
PTYKPKPSYP PTYKSKSSYP SSYKPKKTYP PTYKPKLTYP PIYKPKPSYP PTYKSSYPPR

       730
YKKKISYPSQ Y
```

ⓑ ExPASy Peptide Cutter

| Enzyme | No. of cleavages | Cleavage sites |
|---|---|---|
| **Chymotrypsin** | 86 | 1 7 10 14 22 31 35 39 46 56 62 65 69 72 79 89 93 99 109 119 129 139 143 149 159 169 179 189 199 209 219 229 239 249 259 269 279 289 299 309 319 329 339 349 359 369 375 379 389 399 409 419 423 429 439 449 459 469 475 479 489 499 509 519 529 539 549 559 569 579 589 599 609 619 629 643 653 663 673 683 693 697 703 713 721 731 |
| **Pepsin** | 8 | 1 63 64 68 143 376 423 697 |

Figure S4. a) Amino acid sequence of mgfp-1 from the NCBI data base (Genbank Accession: Q27409) without the predicted N-terminal signal peptide. b) Predicted cleavage sites (ExPASy Peptide Cutter) for mgfp-1 with Chymotrypsin and Pepsin.

**Amphiphilic nature of the coating protein mussel foot protein-1 (mfp-1) from *Mytilus edulis* (mefp-1) and *Mytilus galloprovincialis* (mgfp-1)**

The thread cuticle granules of *M. edulis* and *M. galloprovincialis* both exhibit the characteristic brainy texture described in this paper and in other studies[5]. Since we believe both mefp-1 and mgfp-1 are concentrated in the granules of their respective species[3,4] (Fig. S4), we consider here how specific features of the protein sequence might influence the nanostructural organization of granules observed with TEM and FIB-SEM. Protein sequences from mefp-1 and mgfp-1 can be found in the NCBI database under the GenBank accession numbers AAX23968 and Q27409, respectively. The mefp-1 sequence (AAX23968) possesses a clear signal peptide, but may be missing part of the N-terminal region, since the predicted molecular weight is less than that observed in SDS-PAGE gels[6]. A second published mefp-1 sequence[6], is missing the N-terminal region, but overlaps with AAX23968 and has a high molecular weight – leading us to conclude that the sequence is nearly complete. Regardless, sequence analysis reveals that although mefp-1 (AAX23968) is perhaps slightly truncated, it is extremely homologous to mgfp-1 (Fig. S5a). Both contain a predicted signal peptide cleaved between residue 20 and 21. Both also contain a highly repetitive region consisting of tandem repeats of a Lys and DOPA-rich decapeptide motif that makes up most of the protein length. However, at the N-terminus of both is a largely homologous sequence between 60-80 amino acids that is non-repetitive and markedly less hydrophilic than the repetitive domain. While this cannot be called hydrophobic, it results in the amphiphilic hydropathy profiles for both proteins observed in Fig. S5b-c. We posit that this block co-polymer-like structure facilitates the formation of the bicontinuous brain-like nanostructure of the granules from these species, as show in Fig. 4d.

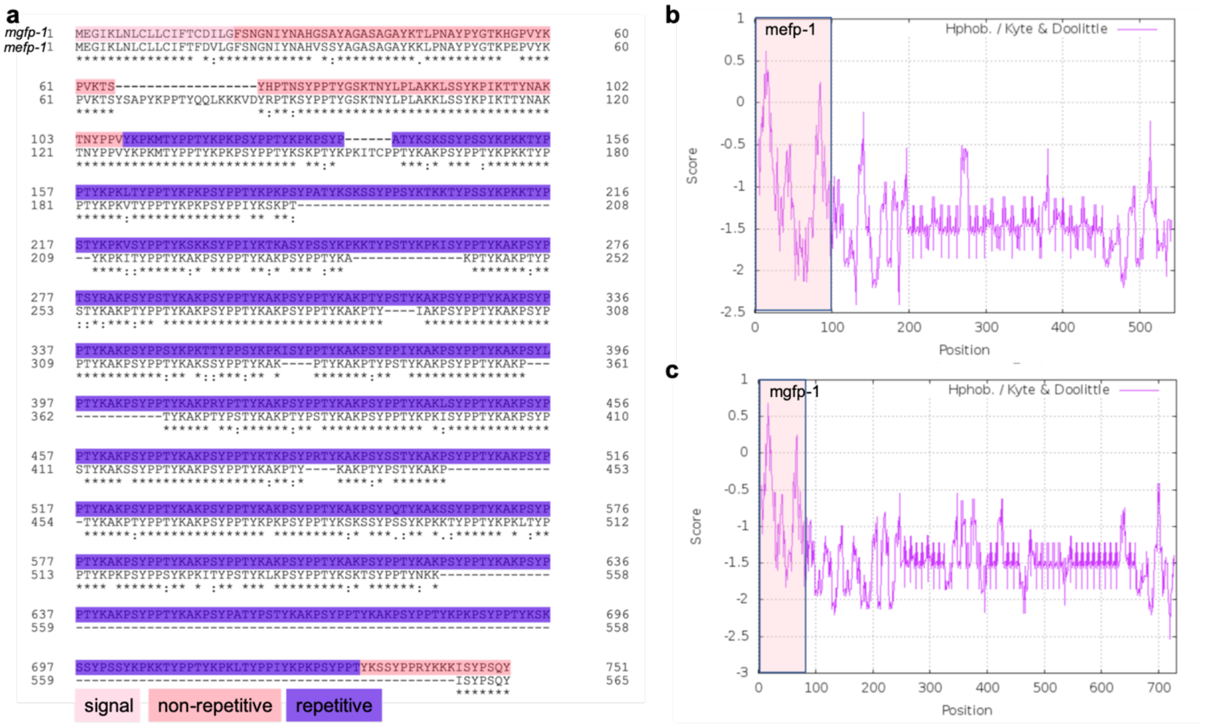


Figure S5. Sequence comparison of mefp-1 and mgfp-1. a) Homology plot of the two sequences reveals a strong similarity. Both consist of a non-repetitive N-terminal domain and an extended repetitive domain consisting of a tandemly repeated decapeptide consensus motif. b and c) hydropathy profiles of (b) mefp-1 and (c) mgfp-1 performed using the Kyte & Doolittle function of ProtScale (EXPASY) with a 9 residues window. The N-terminal non-repetitive domain of both proteins shows a less hydrophilic profile than the repetitive domain.
